# A generalizable method for false-discovery rate estimation in mass spectrometry-based lipidomics

**DOI:** 10.1101/2020.02.18.946483

**Authors:** Grant M. Fujimoto, Jennifer E. Kyle, Joon-Yong Lee, Thomas O. Metz, Samuel H. Payne

## Abstract

Mass spectrometry (MS)-based lipidomics is revolutionizing lipid research with high throughput identification and quantification of hundreds to thousands of lipids with the goal of elucidating lipid metabolism and function. Estimates of statistical confidence in lipid identification are essential for downstream data interpretation in a biological context. In the related field of proteomics, a variety of methods for estimating false-discovery are available, and understanding the statistical confidence of identifications is typically required for data analysis and hypothesis testing. However, there is no current method for estimating the false discovery rate (FDR) or statistical confidence for MS-based lipid identifications. This has slowed the adoption of MS-based lipidomics research, as all identifications require manual inspection and validation to ensure their accuracy. We present here the first generalizable method for FDR estimation, a target/decoy approach, that allows those conducting MS-based lipidomics research to confidently adjust spectral score thresholds to minimize false discovery and to enable full automation of data analysis.

## Introduction

An essential feature of processing and subsequent biological interpretation of high throughput omics data is having confidence in the accuracy of the underlying identifications for the molecule being measured. In mass spectrometry (MS)-based research, this translates to whether one can trust the molecular annotation of a spectrum, i.e. was the assigned spectrum due to the attributed molecule. Statistical confidence of such assignments can be calculated in a number of ways, each with an underlying set of assumptions. To enable high throughput lipidomics data processing and utilization, researchers require a method for estimating statistical confidence of lipid-spectrum matches (LSMs) as they already have for peptide-spectrum matches (PSMs).

For bottom-up MS-based proteomics data, one of the first rigorous methods of estimating the confidence of a peptide-spectrum match was introduced in the PeptideProphet algorithm^1^. In this method, the scores of all PSMs in an LC-MS/MS analysis were plotted and evaluated to identify two overlapping distributions corresponding to the scores of false-positive and true-positive results. After parameterizing the mixed-model distribution^2^, this method could be applied to any LC-MS/MS dataset to identify a score cutoff for a user-specified false-discovery rate. This dramatically improved the quality of research results from bottom-up proteomics data, as users could easily identify high-confidence PSMs and filter out remaining low-quality data, facilitating the automation of data analysis.

Subsequent research for calculating the statistical confidence of PSMs introduced the target/decoy method, where each LC-MS/MS dataset was searched against a database containing both true and intentionally false protein sequences^3^. By always having a set of decoy hits present in search results, the observed true-negative score distribution is trivially used to calculate the false-discovery rate. This is in contrast to the PeptideProphet method which used a model for estimating the false-positive score distribution. The simplicity of the target/decoy method led to rapid adoption in almost all proteomics algorithms and pipelines. Various groups explored how best to create a decoy database, although no single method performed best across diverse experimental designs and organisms^4^. This is in part intuitive, because the purpose of the decoy database is to create a bulk statistic of false identifications. Thus multiple methods for creating false sequences (e.g. reversed or random) are appropriate and perform equally well.

The next advancement in reporting confidence of spectrum identifications came from the generating function approach, which calculates an exact probability of the peptide-spectrum match^5^. Using dynamic programming, the generating function explores the complete sequence space for each and every spectrum. Using this distribution of spectrum-specific scores, the candidate PSM can be accurately evaluated for likelihood. The MSGF algorithm first implemented the generating function^6^, and it has been subsequently adapted and re-implemented by other groups^7,8^.

The chemical differences between lipids and proteins have prevented a straightforward adaptation of either the target/decoy method or the exact probability calculation. The lack of common building blocks analogous to the 20 amino acids has stalled development of both the target/decoy method and a more statistically rigorous generating function. Decoy sequences can be created for proteins by reversing the amino acid sequence or picking a random assortment of amino acids. Although lipids have a building block-like structure, the rigidly defined linkages are not amenable to simple reversing or randomizing. For example, swapping the position of a headgroup and the phosphate linker on a glycerophospholipid, creates a nonsensical molecule. Moreover, there is not one pattern for all lipids, but rather a separate template of linkages and building blocks for each class. These issues have prevented the creation of a target/decoy methodology. Similarly adapting the generating function requires the ability to enumerate all possible lipids to calculate exact statistical significance. Although large libraries have been constructed (LipidBlast^9^ and LipidMaps^10^), these are empirical and not generalizable solutions.

We present here the first report for estimating the false-discovery rate within lipidomics tandem mass spectrometry data using a method for decoy generation. The method generalizes well to any lipid species that has hydrocarbon chains. Thus, while it is not completely universal, it does broadly address the vast majority of lipid categories (i.e., fatty acyls, sphingolipids, glycerophospholipids, and glycerolipids, etc.). Much like the target/decoy approach in proteomics is applied to numerous scoring algorithms (e.g. SEQUEST, Myrimatch, X!Tandem, etc.), this approach can also be applied to other lipid identification software packages.

## Results

### Calculating a probability using runner-up matches

One popular and simple method of calculating PSM probability is embodied in the X!Tandem algorithm^11^. For every spectrum in a global proteomics experiment, there are typically many hundreds of peptide sequences that have a similar precursor mass (e.g. within mass measurement accuracy of the instrument or *m/z* isolation window width used for MS/MS). This set of candidate sequences are scored against the observed fragmentation spectrum, producing a long list of PSMs for a single spectrum. X!Tandem assumes that all non-top-scoring PSMs are false-positive identifications. Using the distribution of these hundreds of false-positive identifications, X!Tandem builds a spectrum-specific null distribution. The probability that the top-scoring peptide is a true-positive is based on this distribution.

We attempted to use an X!Tandem-like method to score LSMs, but were unsuccessful due to the relatively small set of candidate lipids for each precursor mass. We used the LipidMaps database as a source of potential lipid species. Unlike peptides, where one can expect a set of > 500 candidates per spectrum, the list of lipids with similar precursor mass was much smaller (often in the low tens of candidates) and highly varied. This limited set of candidates precluded reliably modeling the false-positive score distribution on a per-spectrum basis (data not shown).

### A generalized decoy molecule for lipids

We introduce a method for creating decoy lipid molecules that is easily generalizable and applies to the vast majority of lipid species and classes. For every lipid in the search space that contains a hydrocarbon chain, we create a decoy lipid that has seven more double bonds per chain than the target chain (see Figure 1). If the target lipid has a hydrocarbon chain with less than 8 carbons, all bonds in the chain become double bonds. Seven double bonds are a very rare event in known lipid species. Indeed, by parsing the LipidMaps database^10^ (download 11/11/2015) we identified only 4 species with 7 double bonds on a carbon chain (LipidMaps identifications: LMFA01030916, LMFA07050066, LMFA07050107, LMPK12140072), out of 40,360 total.

**Figure 1.**
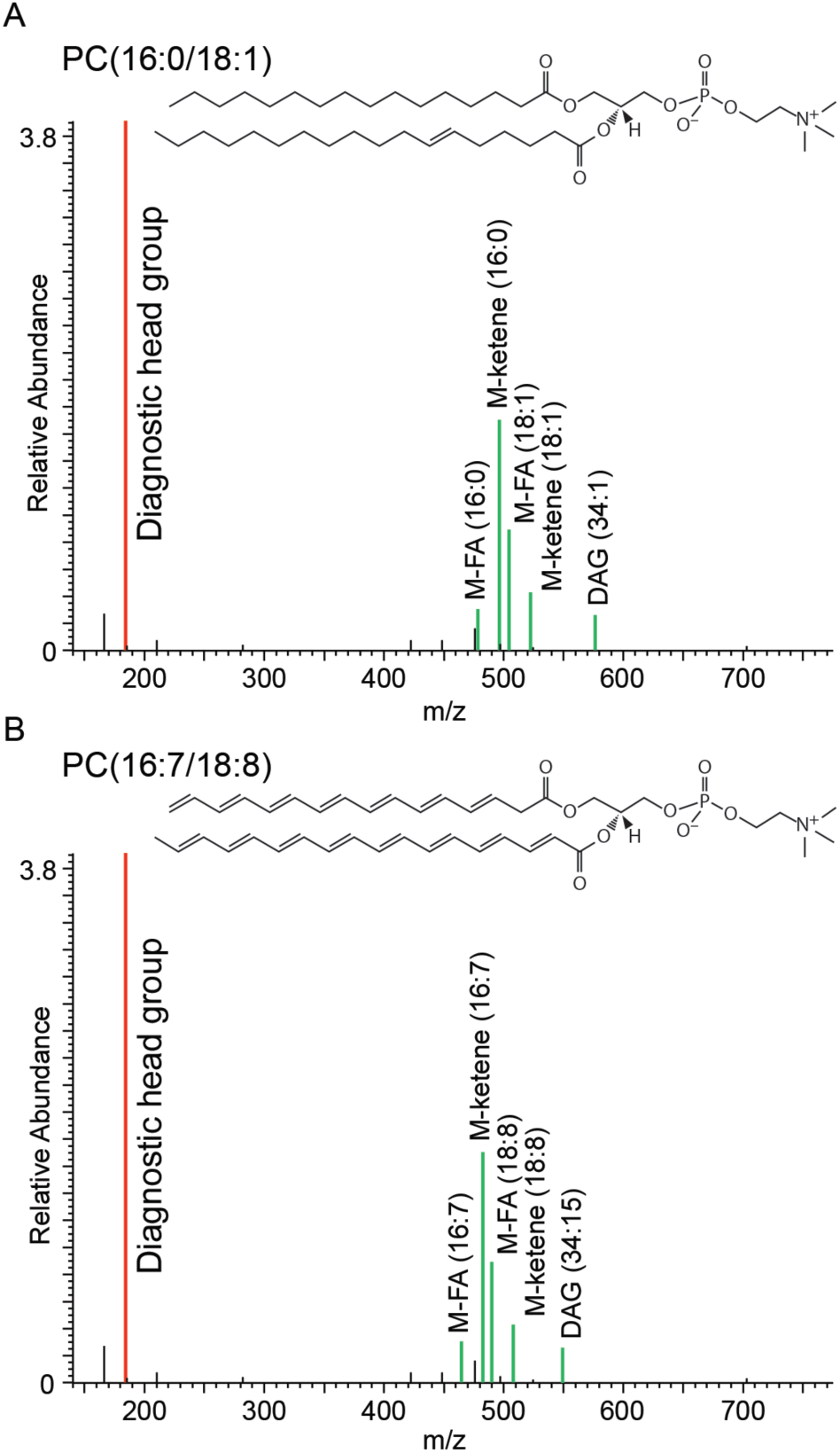
Decoy lipid generation. (A) The structure and MS/MS spectrum of diacylglycerophosphocholine PC(16:0/18:1). The spectrum y-axis is scaled to show the low intensity fragment ions from the acyl tails (green). (B) The structure and hypothetical spectrum of the cognate decoy lipid diacylglycerophosphocholine PC(16:7/18:8). The m/z of acyl-tail fragment ions has been adjusted to account for the loss of mass when making double bonds. The m/z of the diagnostic headgroup ion (red) has not changed as its structure and chemical composition remain unchanged. The intensity scale has been adjusted to show the lower abundance fragment ions.

We chose this option for creating a decoy sequence for the following reasons. First, it is a simple and straightforward implementation that can be applied to a very large variety of lipids. This is essential for the application to global lipidomics research on a wide variety of organisms across all domains of life. Second, this method creates an equivalent number of target and decoy sequences. This is not a requirement for the decoy database methodology, as proteomics sometimes uses a substantially larger set of decoy sequences. For example, the original PeptideProphet methodology used a set of 18 true-positive protein sequences and ∼13,000 decoy sequences. It is, however, convenient to have a 1:1 ratio for target:decoy molecules. Third, the method ensures predictable fragments and the creation of theoretical spectra. Thus any algorithm can easily score decoy lipid molecules. For example, the diacylglycerophosphocholine shown in Figure 1 has a M-ketone (18:1) peak; in the corresponding decoy molecule, this is an M-ketone(18:8) peak and its m/z is adjusted to account for the loss of 14 hydrogen atoms.

### Model Training

It is important to distinguish the goals of creating a reliable method for false-discovery estimation from those for creating an effective LSM scoring function. The goal of a good scoring function is to distinguish true-positive and false-positive identifications. The goal of a FDR estimate method is to report the statistical confidence in a given identification, or more generally to evaluate the success of the scoring function. Explicitly, one can have an accurate method for FDR estimation and have it operate on an imperfect scoring function. This is reminiscent of the early difficulty of proteomics algorithms with phosphoproteomics data^12^ or the lack of confidence in short peptide identifications. The target/decoy approach exposes such shortcomings and provides an opportunity to evaluate possible algorithmic improvements.

We collected true positive and true negative training data from a variety of global LC-MS/MS lipidomics experiments conducted in both positive and negative electrospray ionization, including from virus-infected cell lines, mouse lung tissue, human blood spots, cyanobacteria, and soil. True positive training data came from manual curation and true negative data was taken from all decoy database hits. Through manual curation, we obtained sufficient training data for 23 lipid subclasses: 14 in positive ionization mode and 17 in negative ionization mode (Supplementary Table 1). These training data were inputs to an SVM classifier (see Methods) using four metrics of identification: MS/MS spectrum match score, deviation from expected retention time, isotopic profile match of the precursor ion, and isotopic profile match of the precursor ion -1hydrogen (Supplemental Figure 1). During manual curation of the training data, we noted that the number and type of expected fragment ions varies significantly between different lipid subclasses (as defined by LipidMaps), due to their unique chemistry. Similarly, the same molecule may fragment differently in positive and negative ionization modes. For this reason, we estimate FDR for each subclass and ionization modality separately.

### Algorithm Performance

We examined the utility of the FDR method using a testing dataset comprised of nine LC-MS/MS experiments from human and mouse cell lines. In each lipid subclass, we looked for an overlap between the decoy database hits and low scoring – presumably false-positive – target database hits. In an idealized scenario, target database hits are comprised of a mixed distribution from true positive and false positive matches (Figure 2), as has been previously observed from peptide/spectrum matches^1,3,13^.

**Figure 2.**
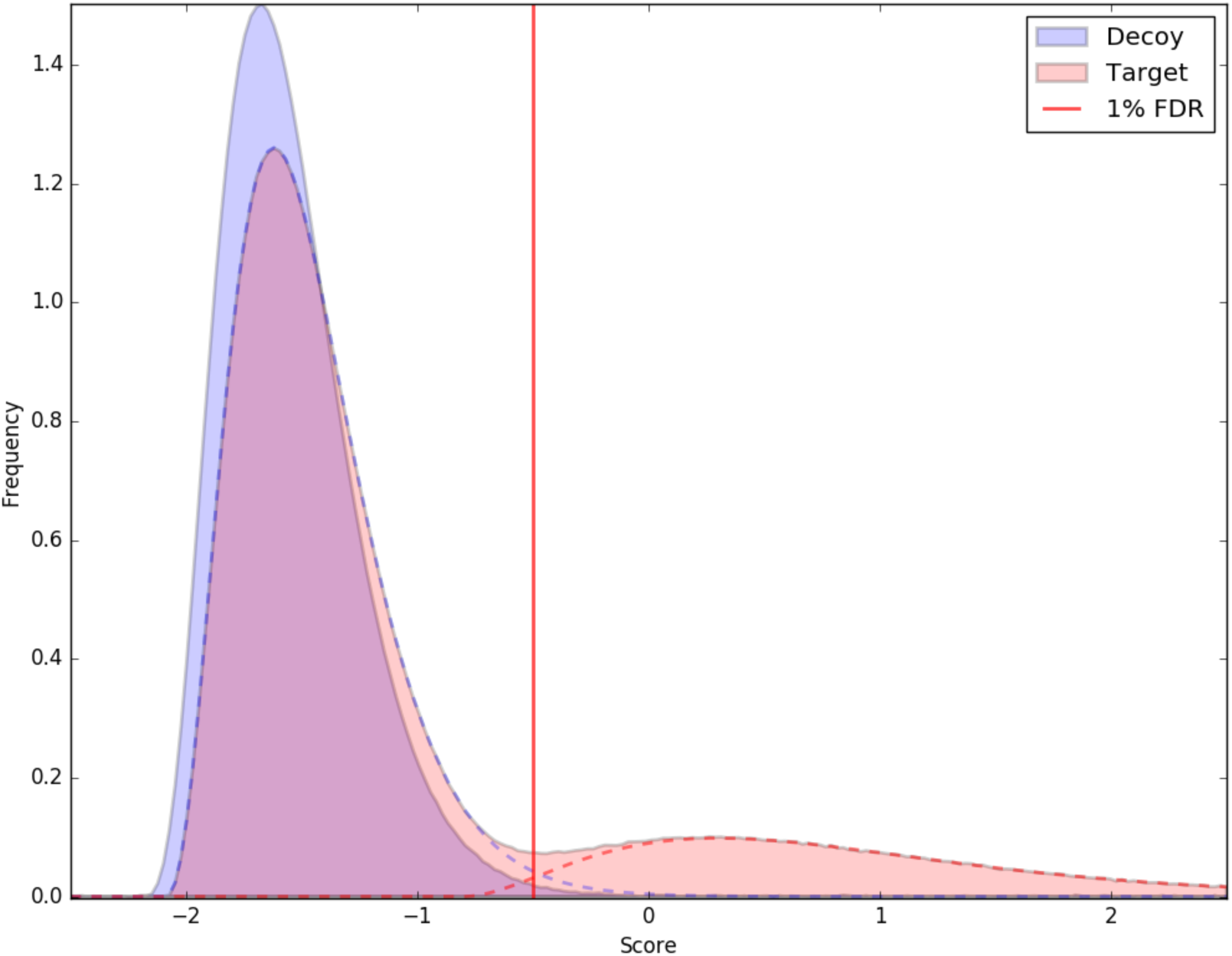
Idealized score distribution of target and decoy hits. The poorly scoring target database hits mimic the true-negative hits to the decoy database. This concept is based on observations in proteomics data, and a similar figure appears in ref ^13^. Dashed lines correspond to the false-positive (blue dash) and true positive (red dash) parts of the target hits score distribution. The red vertical line represents a 1% false discovery rate.

Fourteen of the twenty-three lipid subclasses produced reliable models, including lipids that are biologically important for health and environmental processes, e.g.: ceramides, diacylglycerophosphocholine and triacylglycerols. Some of these models had a more idealized overlap than others. In Figure 3A the diacylglycerophosphoethanolamine subclass (LipidMaps GP0201) has a decoy distribution with a strong peak centered at -1.0 with tails reaching +/- 0.5. Low scoring target database hits mirror this distribution. In Figure 3B, monoacylglycerophosphocholines (LipidMaps GP0105) also have this mirrored overlap, although there are fewer lipid/spectrum matches for this subclass. There are also other subclasses for which the decoy distribution did not as closely mirror low scoring target hits (Figure 3C & D). Although this does not fit the idealized model of target/decoy searches, it still shows how a false (decoy) molecule would be scored in the current scheme. Therefore it is still productive in modeling a false-discovery rate.

**Figure 3.**
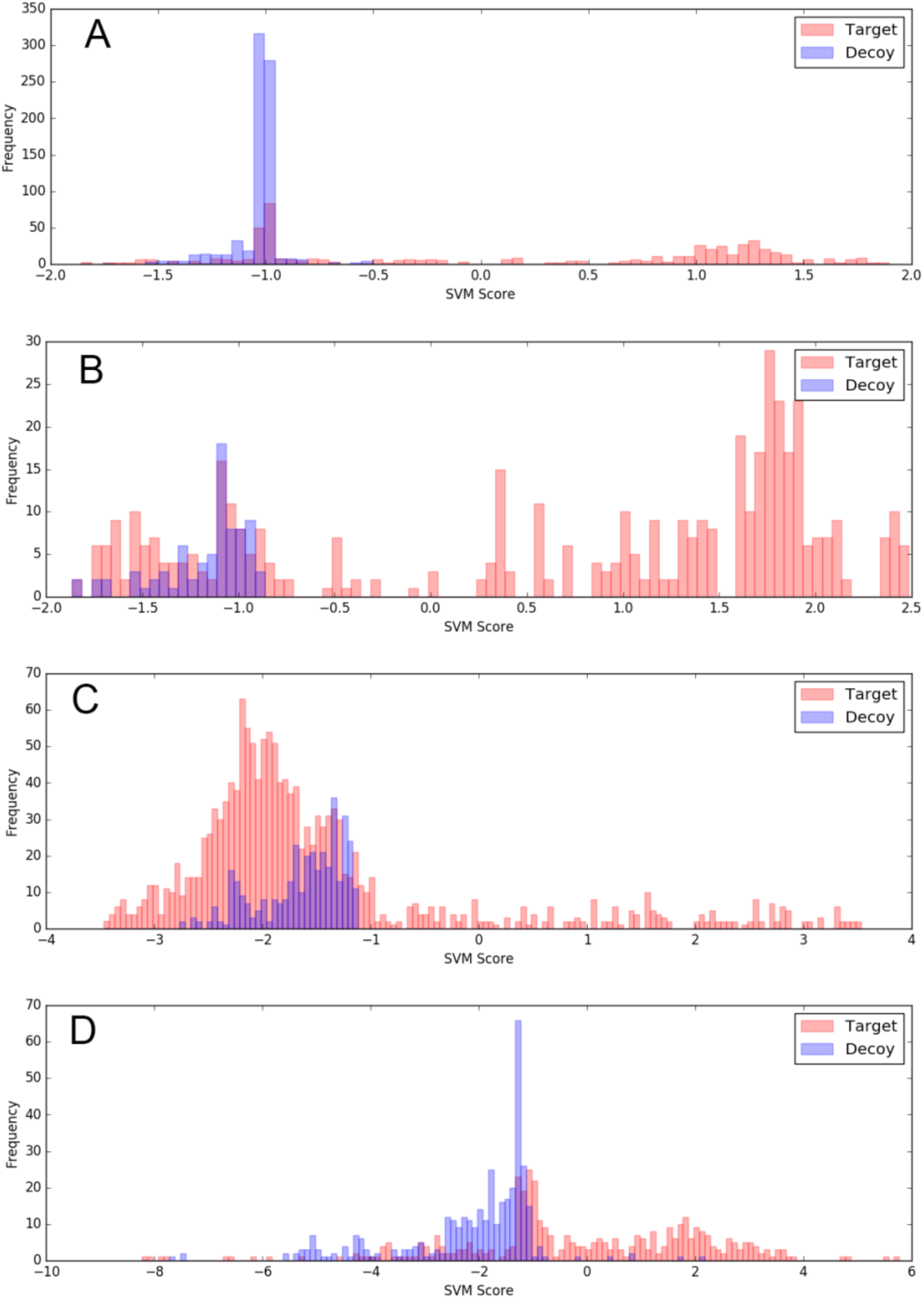
Lipid subclasses with good target/decoy separation. Data is from test set 1, a collection of nine MS/MS lipidomics experiments from human and mouse cell cultures. Although not all subclasses show an idealized distribution of scores, the distribution defines what false identifications would score as, and is separated from high scoring target hits. A. monoacylglycerophosphocholine in positive ionization mode, LipidMaps GP0105 (n=452 target hits, 79 decoy hits). B. diacylglycerophosphoethanolamine in positive ionization mode, LipidMaps GP0201 (n=577 target hits, 748 decoy hits). C. diacylglycerophosphoserine in positive ionization mode, LipidMaps GP0301 (n=1454 target hits, 360 decoy hits). D. ceramide phosphocholine (sphingomyelins) in positive ionization mode, LipidMaps SP0301 (n=412 target hits, 374 decoy hits).

For some lipid classes, the separation between target and decoy scores was not resolved because there were too few decoy hits or the hits are disperse and there is no clear separation between target and decoy (Figure 4A & B). Six subclasses fell into this category. For example, diacylglycerophosphoinositols have hundreds of high scoring spectra in the target database, but few decoy hits; with only few low scoring true-negatives it may not be appropriate to assign a statistical confidence associated with these models. Although the current FDR implementation will report a value based on these distributions, they should be used with caution. Which specific lipid subclasses fall into this category will change with each experiment as the number and type of lipids is expected to be distinct.

**Figure 4.**
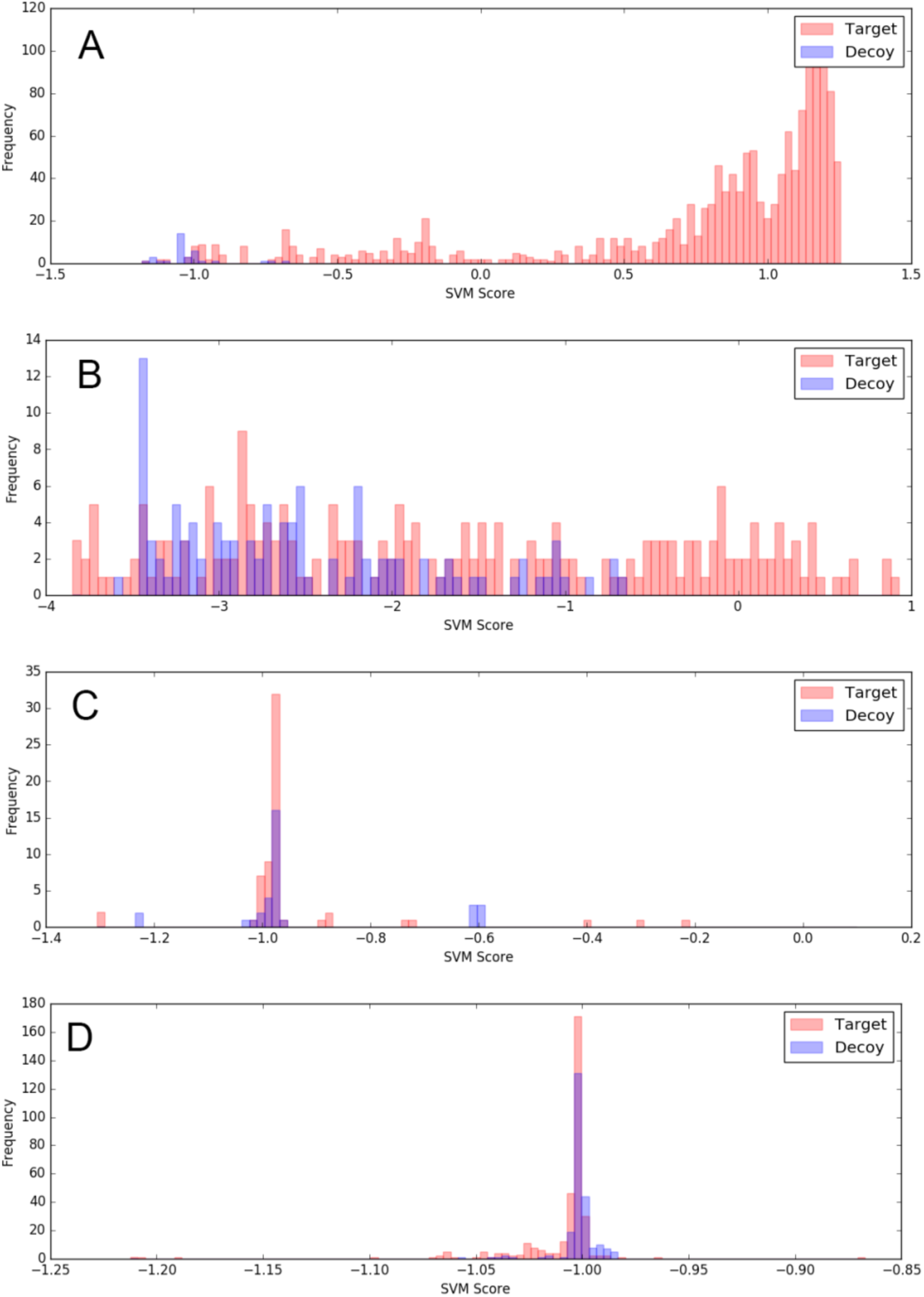
Lipid subclasses with poor target/decoy separation. Data is from test set 1, a collection of nine MS/MS data files from human and mouse cell cultures. In A, although there are many high scoring target hits, the number of decoy hits is too small to reliably model the score distribution. In B, the decoy score distribution is disperse and poorly sampled, which makes it difficult to model the underlying distribution. In C and D, the target and decoy distributions show clear overlap, but there are no good scoring target hits. A. diacylglycerophosphoinositols in negative ionization mode, LipidMaps GP0601 (n=1464 target hits, 34 decoy hits). B. monoacylglycerophosphoethanolamines in positive ionization mode, LipidMaps GP0205 (n=226 target hits, 112 decoy hits). C. monoacylglycerols in positive ionization mode, LipidMaps GL0101 (n=61 positive hits, 33 decoy hits). D. glycosyldiacylglycerol in positive ionization mode, LipidMaps GL0501 (n=334 target hits, 237 decoy hits).

A final challenge is lipid subclasses whose fragmentation pattern is difficult to accurately score. Monoacylglycerols (LipidMaps GL0101) are comprised simply of a fatty acid and a glycerol backbone and only reliably produce two fragment ions in positive ionization. The paucity of fragments generated by CID makes it difficult for a scoring function to discriminate between true and false spectral identifications. Testing data for this subclass produced nearly identical score distributions for the target and decoy hits (Figure 4C). Another example of challenging scoring is when a subclass groups together molecules that behave differently in MS/MS fragmentation, like glycosyldiacylglycerol, GL0501 at LipidMaps. This subclass contains three types of lipids with very distinct diagnostic ions and fragmentation patterns: MGDG, DGDG, and SQDG. Thus the score of true positive lipid-spectrum matches will be an amalgam of distributions. When applied to the machine learning SVM approach this results in a poor separation for target and decoy hits (Figure 4D).

### Consistency of decoy distribution

As some of the lipid subclasses have decoy distributions that did not match the idealized scenario depicted in Figure 2, we attempted to determine whether the distribution of decoy scores for a given lipid subclass was consistent across different experimental samples. Therefore, we applied the target/decoy method to a second test data set from mouse lung tissue (9 positive and 9 negative ionization MS/MS files). For each of the 23 subclasses, the decoy distribution from the original test set 1 was compared to the decoy distribution from test set 2. Figure 5 shows several such comparisons in detail. Although the number of observations for each decoy distribution varies between test set 1 and 2 (as expected), the score distributions were consistent.

**Figure 5.**
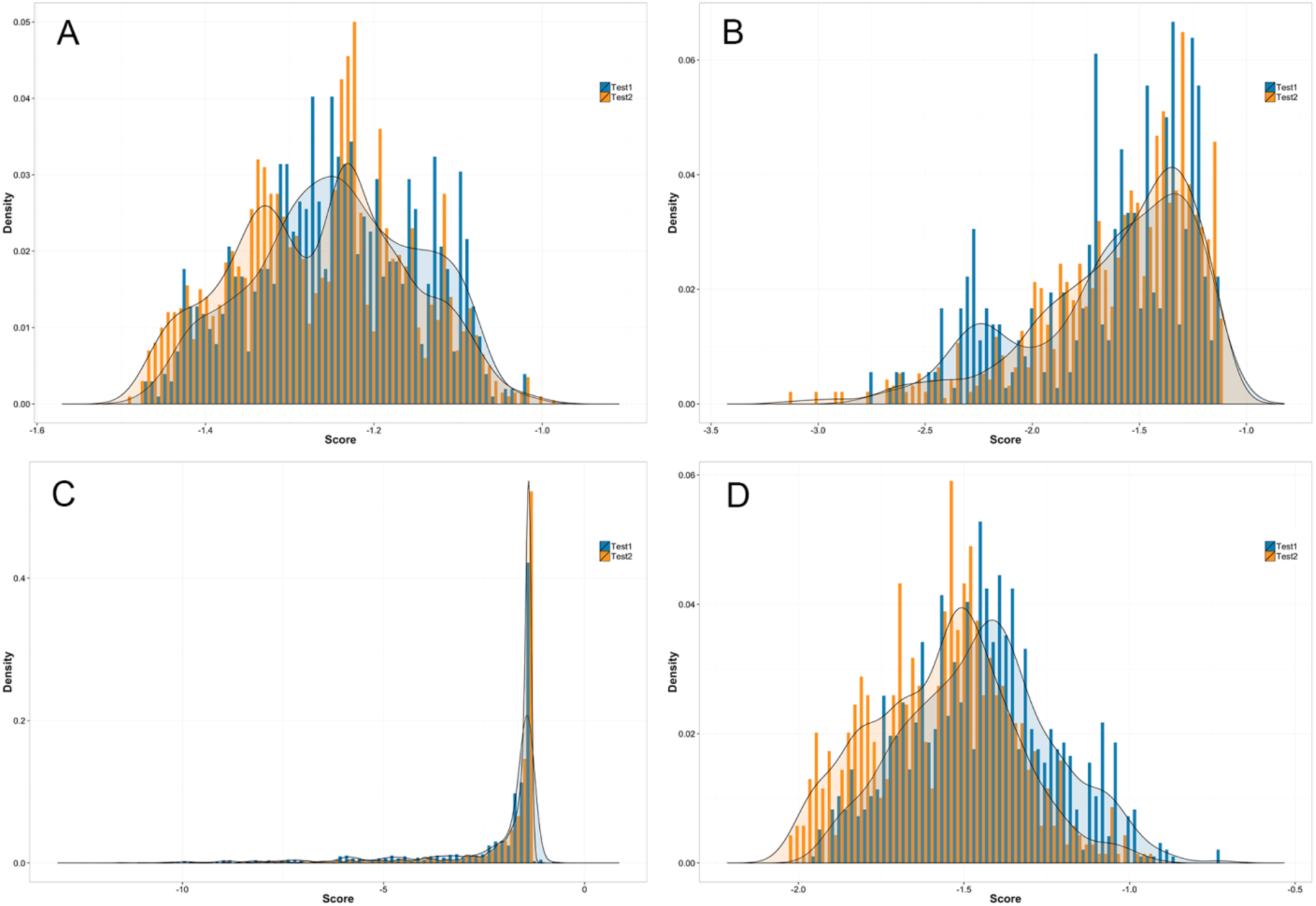
Consistency of decoy distribution. The decoy score distribution for the two different test sets is plotted against each other. Plotted in blue are decoy hits from test set 1 (nine experiments from human and mouse cell cultures); plotted in orange are decoy hits from test set 2 (nine experiments from mouse lung tissue). A. N-acylsphinganines (dihydroceramides) in positive mode ionization, LipidMaps SP0202 (n=1019 from test set 1, 1999 from test set 2); B. diacylglycerophosphoserines in positive ionization mode, LipidMaps GP0301 (n=360 for test set 1, 940 for test set 2); C. diacylglycerols in positive ionization mode, LipidMaps GL0201 (n=1053 for test set 1, 2268 for test set 2); D. N-acylsphingosines (ceramides) in negative ionization mode, LipidMaps SP0201(n=966 for test set 1, 694 for test set 2).

## Discussion

The proteomics community has benefited from algorithms such as SEQUEST, X!Tandem, MSGF and others for 20 years, allowing researchers to rapidly identify peptides and their corresponding proteins. Equally important have been methods for assessing the false discovery rate, which allows researchers to understand the statistical confidence of peptide and protein identifications. The ability to accurately measure and report confidence was necessary for the widespread adoption of proteomics data in environmental and biomedical research.

In the related fields of metabolomics and lipidomics, a method for assessing the false discovery rate of spectral matches has not been established. The current gold standard for confident identification of metabolites is to match data (e.g. tandem mass spectra, retention time, accurate mass) from experimental measurements to reference libraries obtained from analyses of purified chemical standards, an approach endorsed by the Metabolomics Standards Initiative of the Metabolomics Society^14^. Confidence in the identifications is typically asserted via manual inspection of the spectral match, or a non-statistical score cutoff. This approach is untenable as metabolomics and lipidomics researchers increasingly try to manage tens to hundreds of gigabytes of data per experiment. Further, the criteria for confidently identifying metabolites and lipids is evolving, and the existing criteria have not yet been broadly adopted^15,16^. The development of a method for robustly estimating FDR, regardless of the approach used for metabolite or lipid identification, could facilitate the establishment of consensus criteria by the research community^17^.

The target/decoy method presented here is a simple option for the false discovery rate of lipid-spectrum matches. For many biologically significant classes of lipids, this method works well. For the classes where scoring methods currently fall short, the false discovery rate calculations provide a platform for objective comparison of scoring models. Potential improvements include more advanced algorithms and instrumentation/fragmentation techniques to increase the number and intensity of reliable lipid fragment ions (e.g. MS^3^). We feel that the FDR estimation method described here will be invaluable in guiding the next stages of lipidomics research. For proteomics, introducing an FDR estimation method ushered in an era of intense computational interest and improvement, with dozens of new software tools being released. New instrumentation was also developed to address the need for more consistent fragmentation or better mass measurement. We hope and anticipate that a similar surge will now form around lipid-spectrum identification.

## Acknowledgments

We thank Drs. Charles K. Ansong, Erin S. Baker, Janet K. Jansson, and Malak M. Tfaily at PNNL and Amie J. Eisfeld and Yoshihiro Kawaoka at the University of Wisconsin for the samples used to generate the lipidomics data needed for this work. Funding was provided by the Laboratory Directed Research and Development Program (LDRD) at Pacific Northwest National Laboratory (PNNL), the U.S. Department of Energy (DOE), Office of Science, Office of Biological and Environmental Research (BER) under the PNNL Pan-omics program, and the National Institutes of Health, National Institute of Allergy and Infectious Diseases grant U19AI106772, National Heart, Lung, and Blood Institute grant U01HL122703. Lipidomics measurements to generate data for development of the FDR approach were performed in the Environmental Molecular Sciences Laboratory, a national scientific user facility sponsored by the U.S. DOE BER and located at PNNL in Richland, Washington. PNNL is a multi-program national laboratory operated by Battelle for the DOE under Contract DE-AC05-76RLO 1830.

## Author Contribution

Study conception and design: JEK, TOM, SHP. Computation and statistical analysis: GMF, JYL, SHP. Data interpretation: GMF, JEK, JYL. Manuscript writing: all authors.

The authors declare no competing financial interests.

## Methods

### Datasets used

Lipidomics data from 8 different experiments comprising 79 LC-MS/MS analyses were included in training data in this study. Eight analyses were of total lipid extracts derived from human blood spots and serum, 25 of virus and mock infected human epithelial cell lines (Calu-3 purchased from ATCC http://www.atcc.org/products/all/HTB-55.aspx), 32 of virus and mock infected mouse lung tissue, 4 of mouse lung tissue, 6 from two different soil types, and 4 from cyanobacteria cultures. All mice were female; all animal work was approved by the Institutional Animal Care and Use Committee of Pacific Northwest National Laboratory prior to initiation of the study. Test set one consists of nine LC-MS/MS analyses in both positive and negative mode; 6 samples were from Calu-3 cells, and three samples were from primary cells derived from mouse lymph nodes. Test set two consists of nine LC-MS/MS analyses in both positive and negative modes; all nine samples were from mouse lung tissue. For each of these samples we used the following extraction and analysis protocol. A Waters NanoAquity UPLC system (Waters Corporation, Milford, MA) interfaced with a Velos-ETD Orbitrap mass spectrometer (Thermo Scientific, San Jose, CA) was used for LC-ESI-MS/MS analyses. Total lipid extracts (obtained using a modified Folch extraction^18^) were reconstituted in MeOH and injected onto a C18 reversed-phase column (HSS T3. 1.0 mm × 150 mm × 1.8 µm particle size; Waters). Lipids were separated over a 90 min gradient elution (mobile phase A: ACN/H_2_O (40:60) containing 10 mM ammonium acetate; mobile phase B: ACN/IPA (10:90) containing 10 mM ammonium acetate) at a flow rate of 30 µl/min. Samples were analyzed in both positive and negative ionization using HCD (higher-energy collision dissociation) and CID (collision-induced dissociation) to obtain high coverage of the lipidome. The raw mass spectrometry datasets have been uploaded to the MassIVE repository (http://massive.ucsd.edu), under submission MSV000079770.

### Method for Decoy Generation

The target lipid database was obtained from LipidMaps. Lipid decoys are based on these lipid species with seven double bonds added to each of the lipid acyl chains. For example, if the target species was a PC(16:1/16:0), then the decoy would be PC (16:8/16:7). Because MS/MS methods are not typically able to localize double bonds within the hydrocarbon chain, the structure of this decoy molecule is not specified beyond the number of double bonds. If the acyl chain contained fewer than eight carbons, all carbons in the chain would be double bonded. For example, a fully saturated hydrocarbon chain of carbon length seven, commonly annotated as 7:0, would become a decoy with six double bonds, annotated as 7:6. This method is appealing because it is generalizable to any class of lipid that has a hydrocarbon chain, e.g. PC, DG, TAG, etc. The decoy database file used in this manuscript is available as Supplemental Table 2 and 3.

### Software implementation of the FDR

A support vector machine was used to create a single score from the four feature metrics of lipid-spectrum matches. The model was trained using 41 positive ionization and 37 negative ionization datasets from experiments with environmental and mammalian cells. Datasets were first run through LIQUID for initial lipid identification and scoring. True positive data points were derived from manually curated lipid/spectrum identifications (n=9251). True negative data points were derived from all hits to the decoy database (n=214737). See Supplemental Table 1 for a complete list of the training data. These training data were run through the SVM training script written in Python and using scikit-learn machine learning library. (The software for training the SVM models and also for using the models are available at http://github.com/PNNL-Comp-Mass-Spec/LipidFDR) The model trained on four data quality metrics from the LIQUID output: retention time, cosine similarity of the observed precursor isotope envelope to the theoretical isotope, cosine similarity of the observed precursor isotope envelope to the M-1 theoretical envelope, and LIQUID spectrum score. Because the LIQUID spectrum score is not uniform across different lipid classes (due to different numbers of fragment ions generated in MS/MS), models were created for every class of lipid and both positive and negative ionization mode. For SVM models, a radial basis function (RBF) kernel was employed and random permutations cross validation was used to optimize the kernel parameters.

To test the models we used datasets which are unrelated to those used for training. The testing datasets were processed by LIQUID for initial identification and scoring. For the target database, we used a HCD tolerance of 30 PPM as well as identifications to one per MS/MS spectra. For the decoy database, we used an HCD tolerance of 50 PPM to account for small mass shifts difference expected between true and decoy lipid species as well as allowing any number of decoy identifications per MS/MS spectra. From both target and decoy searches, the SVM classifier scored identifications based on the distance to a separating hyperplane of a trained model to classify the curated lipid/spectrum pairs from the true negative. Scores from target and decoy identifications were transformed into probability density functions, and the FDR estimate was calculated based on the ratio of these two distributions at a given score:

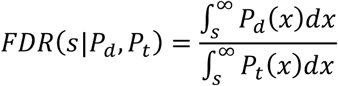

where, *P*_*d*_(*s*) and *P*_*t*_(*s*) indicate probability distributions for decoy scores and for target scores, respectively.

## Figures

**Supplementary Figure 1.**
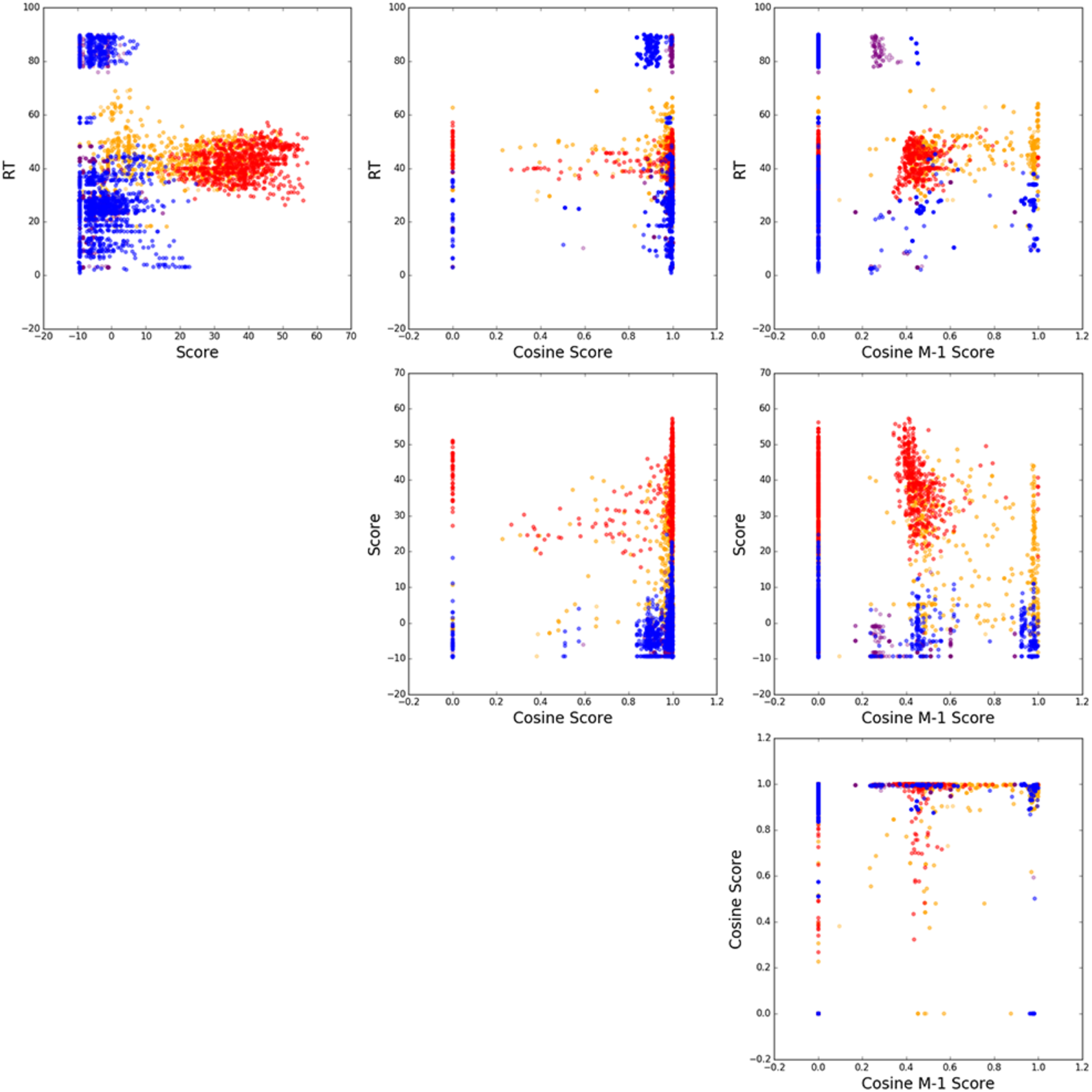
Metrics used to create the SVM score. Data is shown for training and testing lipid/spectrum matches. Red is for true positive training data (n = 866); blue is for true negative training data (n = 3389). Orange is for all target hits on testing data (n = 1421); purple is for all decoy hits on testing data (n = 817). This data is from lipid/spectrum matches to diacylglycerophosphoglycerols in negative ionization mode (LipidMaps GP0401). Similar charts were created for all subclasses as a way to visualize the effective separation of target and decoy hits by the various score metrics.

